# Co-immunoprecipitation with MYR1 identifies three additional proteins within the *Toxoplasma* parasitophorous vacuole required for translocation of dense granule effectors into host cells

**DOI:** 10.1101/867788

**Authors:** Alicja M. Cygan, Terence C. Theisen, Alma G. Mendoza, Nicole D. Marino, Michael W. Panas, John C. Boothroyd

## Abstract

*Toxoplasma gondii* is a ubiquitous, intracellular protozoan that extensively modifies infected host cells through secreted effector proteins. Many such effectors must be translocated across the parasitophorous vacuole (PV) in which the parasites replicate, ultimately ending up in the host cytosol or nucleus. This translocation has previously been shown to be dependent on five parasite proteins: MYR1, MYR2, MYR3, ROP17, and ASP5. We report here the identification of several MYR1-interacting and novel PV-localized proteins via affinity purification of MYR1, including TGGT1_211460 (dubbed MYR4), TGGT1_204340 (dubbed GRA54) and TGGT1_270320 (PPM3C). Further, we show that three of the MYR1-interacting proteins, GRA44, GRA45, and MYR4, are essential for the translocation of the *Toxoplasma* effector protein GRA16, and for the upregulation of human c-Myc and cyclin E1 in infected cells. GRA44 and GRA45 contain ASP5-processing motifs, but like MYR1, processing at these sites appears to be nonessential for their role in protein translocation. These results expand our understanding of the mechanism of effector translocation in *Toxoplasma* and indicate that the process is highly complex and dependent on at least eight discrete proteins.

**Importance:** *Toxoplasma* is an extremely successful intracellular parasite and important human pathogen. Upon infection of a new cell, *Toxoplasma* establishes a replicative vacuole and translocates parasite effectors across this vacuole to function from the host cytosol and nucleus. These effectors play a key role in parasite virulence. The work reported here newly identifies three parasite proteins that are necessary for protein translocation into the host cell. These results significantly increase our knowledge of the molecular players involved in protein translocation in *Toxoplasma*-infected cells, and provide additional potential drug targets.

## Introduction

*Toxoplasma gondii* is an obligate intracellular parasite that can cause severe illness in immunocompromised individuals and the developing fetus. It is estimated to infect up to a third of the world’s population, and has an unparalleled host range, infecting virtually any nucleated cell in almost any warm-blooded animal (1). In order to survive within a host cell, *Toxoplasma* tachyzoites, the rapidly-dividing, asexual stage of the parasite, establish a replicative niche, the parasitophorous vacuole (PV), whose membrane (PVM) acts as the interface between parasite and host. While the PV protects intracellular *Toxoplasma* from clearance by the innate immune system, it also acts as a barrier that *Toxoplasma* must overcome in order to hijack host resources.

*Toxoplasma* extensively modifies the host cells it infects via secreted effectors, either rhoptry (ROP) or dense granule (GRA) proteins, which it introduces into the host during or following invasion (2). In recent years, several *Toxoplasma* GRAs, including GRA16, GRA24, IST, HCE1/TEEGR, GRA28, and GRA18, have been identified that are translocated across the PVM into the host cell cytosol and/or nucleus, where they can have profound effects on host processes (3–9). The machinery that is responsible for the translocation of these effectors across the *Toxoplasma* PVM is incompletely defined. A recent forward genetic screen identified several parasite proteins essential for GRA protein translocation, including MYR1, MYR2, MYR3, (named for their effect on host c-<u>My</u>c regulation) and the rhoptry-derived protein kinase, ROP17 (10–12). Precisely how these proteins function to promote protein translocation across the PVM is poorly understood. Of the four, the only protein with a known biochemical function is ROP17, a serine/threonine protein kinase that phosphorylates host, and perhaps parasite proteins at the PVM (6, 13, 14).

In addition to MYR1, MYR2, MYR3, and ROP17, an active *Toxoplasma* aspartyl protease V (ASP5), which proteolytically processes secreted proteins at the amino acid sequence “RRL” (also known as a *<u>T</u>oxoplasma* <u>ex</u>port <u>el</u>ement, or TEXEL), is also required for the translocation of all exported GRAs studied thus far (5, 6, 8, 9, 15–17). In *Plasmodium*, the homolog of ASP5, plasmepsin V, appears to “license” many proteins for export across the PVM by proteolytically processing them at a *Plasmodium* export element (“RxLxE/Q/D”) (18–21). Intriguingly, and as for *Plasmodium* (22, 23), not all of *Toxoplasma’s* exported GRAs contain “RRL” motifs (e.g. GRA24, GRA28, and HCE1/TEEGR lack such an element), which leaves open the possibility that ASP5’s role in translocation is in processing the translocation machinery, rather than the effectors themselves. Indeed, MYR1 is processed by ASP5, but this processing is not necessary for protein export, as unprocessed full length MYR1 harboring a mutated “RRL” motif can still promote the translocation of the effector GRA24 to the host nucleus (24). The role of ASP5 processing of MYR1, therefore, remains unknown.

To learn more about the mechanism of protein translocation in *Toxoplasma*, and to complement the genetic approaches taken previously, we report here the use of MYR1 as “bait” for immunoprecipitation followed by mass spectrometry (IP-MS) to identify putative MYR1-associated proteins that are involved in effector translocation. Of the many associating proteins, at least eleven are shown here or were previously known to be PV-localized and, of these, three additional proteins are now shown to be required for GRA translocation across the PVM. Interestingly, all three of these new components contain “RRL” motifs, with two confirmed to be cleaved in an ASP5-depndent manner; yet, like MYR1, cleavage at these sites appears not to be required for their translocation function. Thus, we have expanded the list of proteins involved in GRA translocation to eight while also expanding the enigma of why at least three of these components are proteolytically processed without any apparent impact on their one known function.

## Results

We previously reported the use of a forward genetic screen to identify *Toxoplasma* genes required for the induction of human c-Myc. This identified *MYR1*, *MYR2*, *MYR3*, and *ROP17* as essential for the translocation of effector proteins across the PVM (10–12). Two of these proteins, MYR1 and MYR3, were found to co-precipitate with each other (11), and we hypothesized that MYR1 functions in complex with other yet unidentified proteins to facilitate effector translocation across the PVM. Given the small but significant reduction in plaque size observed when growing strains deleted in MYR1, MYR2, and MYR3 on human foreskin fibroblasts (HFFs) (11), we also reasoned that the genetic approach might also miss genes whose disruption substantially reduces fitness.

To identify additional MYR1-associating proteins, therefore, we adopted a biochemical approach. Specifically, we immunoprecipitated 3xHA-tagged MYR1 from HFFs infected for 24 hours with an RH::*MYR1-3xHA* strain, or from an untagged RH strain to control for proteins that co-precipitate with the anti-HA beads nonspecifically (**Fig. 1A**). Liquid chromatography-tandem mass spectrometry (LC-MS/MS) was performed on the eluates and the identified parasite proteins were ranked by the ratio of average normalized spectral abundance factors (NSAFs) for a given protein in the RH:*MYR1-3xHA* lysates compared to the RH control (25). This mass spectrometry experiment was performed twice (IP 1 and IP 2). As expected, MYR1 was the most enriched protein in both biological replicates (**Fig. 1B**). Additionally, several PV- or PVM-localized GRA proteins were highly enriched (enrichment score >10) in the MYR1-3xHA immunoprecipitations over the untagged RH control, including GRA44, CST1, GRA52, MAG1, PPM11C, GRA50, MAF1a, GRA7, and a GRA12 paralog, in addition to two exported effector proteins, GRA16 and GRA28, of which GRA16 has been shown to be exported in a MYR1-dependent manner (10) (**Fig. 1B, File S1**). The large number of enriched PV- and PVM-localized proteins may be explained by the mild detergent conditions used (0.1% NP-40), which were chosen in an attempt to maintain associating proteins, although these proteins might also be associating with one another in large, non-specific complexes or lipid rafts (26).

**Figure 1.**
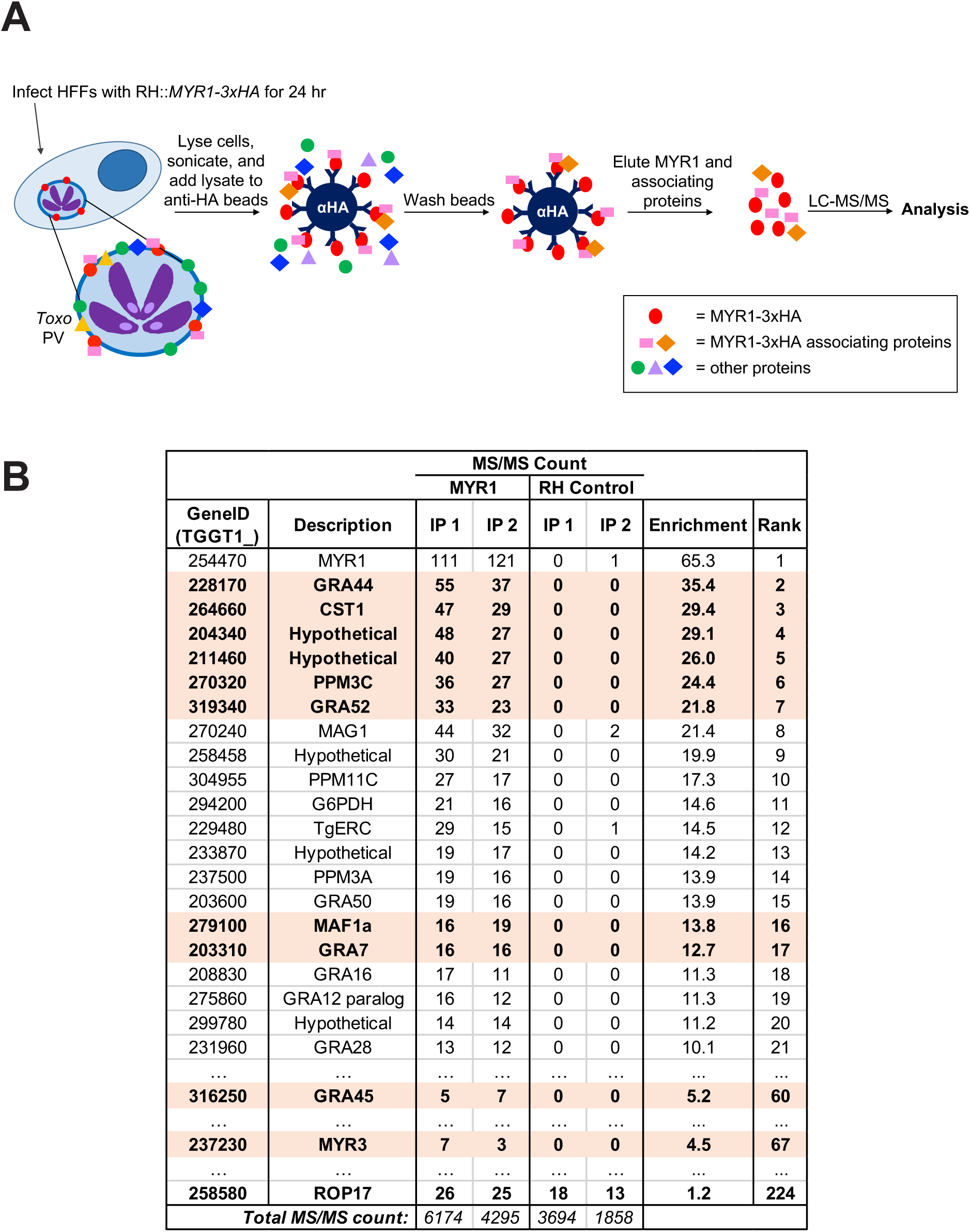
MYR1-3xHA immunoprecipitation identifies many MYR1-associating *Toxoplasma* proteins. A. Schematic of MYR1 IP-MS workflow. B. Results of IP-MS analysis. Mass spectrometry was performed on immunoprecipitated material as depicted in Fig. 1A and the number of spectral counts was determined for all identified proteins. This experiment was performed twice (IP 1 and IP 2) for both RH::MYR1-3xHA and an RHΔ*hpt* untagged control. The identified *Toxoplasma* proteins from the two experiments were ranked according to the average NSAF enrichment in the MYR1-3xHA-expressing strain relative to the untagged RH control after adding a nominal single count to all results, enabling a ratio to be determined (Enrichment and Rank). The full dataset, including associating host proteins, is presented in File S1. Displayed here are the majority *Toxoplasma* protein identifiers (TGGT1_), i.e., the proteins that contain at least half of the peptides belonging to a group of proteins that cannot be unambiguously identified by unique peptides, the descriptive name for each protein (Description), and the corresponding number of spectral counts detected (MS/MS count) for all *Toxoplasma* proteins with an average enrichment score greater than 10. Also shown are data for the proteins GRA45, MYR3, and ROP17. Genes chosen for subsequent disruption are highlighted in orange.

Importantly, and also as expected, the known MYR1-associating protein, MYR3, was enriched in the MYR1-3xHA immunoprecipitations, albeit with an enrichment score (4.5) that did not put it in the top 20 most enriched proteins (**Fig. 1B, File S1**). Of note, ROP17 was not substantially enriched (enrichment score = 1.2) and no peptides for MYR2 were detected, but neither protein has previously been found to associate with MYR1 and so this was not unexpected. Human proteins with an enrichment score >10 include Filamin-C (FLNC), DNA-dependent protein kinase catalytic subunit (PRKDC), sarcoplasmic/endoplasmic reticulum calcium ATPase 2 (ATP2A2), and Alpha-N-acetylglucosaminidase (NAGLU) (**File S1**). As our focus was on parasite proteins only, the potential role of these human proteins in *Toxoplasma* infection was not further investigated.

To screen for a possible role in GRA effector translocation, we focused on the top 6 most enriched parasite proteins: GRA44 (TGGT1_228170), CST1 (TGGT1_264660), TGGT1_204340, TGGT1_211460, PPM3C (TGGT1_270320), and GRA52 (TGGT1_319340). GRA45 (TGGT1_316250) was also pursued because it is a known binding partner of the top hit, GRA44 (27), and it also had a substantial enrichment score of 5.2 in the immunoprecipitations (**Fig. 1B**). Interestingly, two well-characterized PV proteins that have not previously been described to be involved in effector translocation, GRA7 (TGGT1_203310) and MAF1a (TGGT1_279100), were substantially enriched and, since antibodies and gene knockouts for both were readily available, we included these in the list of genes to explore further. Lastly, as a positive control for a protein whose disruption is known to prevent effector translocation, we also included MYR3 in the pipeline for gene disruption and testing. All ten proteins chosen for further analysis are highlighted in orange in Fig. 1B.

With the exception of ASP5, and as might be expected, all proteins so far published as required for effector translocation across the PVM localize to the PV/PVM (2). Of the ten proteins chosen for further analysis, GRA44, GRA45, CST1, GRA7, GRA52, and MAF1a, are all known to be PV/PVM-localized (27–31). The localization of 211460 and PPM3C has not been reported, but both include predicted signal peptides (see below) as does 204340 which has been described as possibly micronemal (32). We therefore set out to localize these three proteins within infected cells. To do this, we generated populations of parasites in which each of the three genes was endogenously modified to encode a 3xHA-tag immediately before the stop codon and then assessed the protein’s localization by immunofluorescence assay (IFA). Correct integration of the 3xHA-tag into the appropriate locus was confirmed by PCR and by checking for an appropriately sized HA-tagged protein via western blotting. The results (**Fig. 2A**) showed major bands at ∼130 kDa, ∼110 kDa, and ∼70 kDa for 211460, 204340, and PPM3C, respectively. In the case of 204340 and PPM3C, this is close to the predicted sizes of ∼97 kDa and ∼60kDa (ToxoDB v45). For 211460, however, the mobility is significantly retarded relative to its predicted size of ∼100 kDa. This could be due to its acidic pI of 4.91 (ToxoDB v45) which is known to reduce protein mobility on SDS-PAGE (33), and/or to post-translational modifications (all three proteins are reported to be phosphorylated (31; ToxoDB v45)). This same slower-than-expected mobility for the major band was seen for an independently generated, cloned line expressing HA-tagged 211460 (**Fig. S1A**) and so we conclude that this is the correct mobility for this protein. Interestingly, both the 211460-3xHA tagged population and single clone also showed a smaller but considerably weaker band at around the expected size (∼100 kDa). Whether this smaller MYR4 product is biologically relevant, or is simply a product of protein degradation, is unclear.

**Figure 2.**
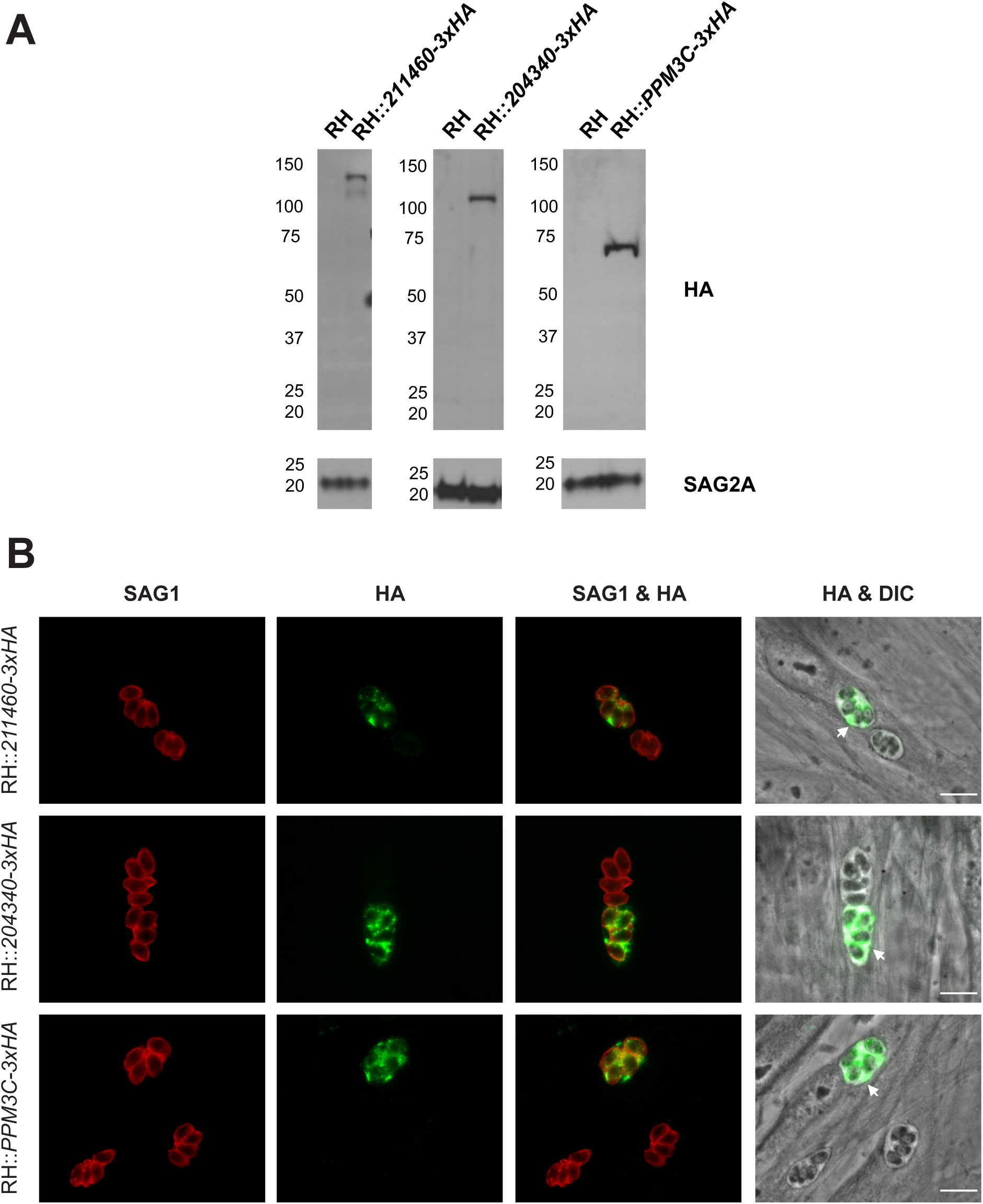
211460, 204340, and PPM3C localize to the *Toxoplasma* parasitophorous vacuole in infected cells. A. Western blot of endogenously tagged parasite proteins. HFFs were infected with RHΔ*hpt*Δ*ku80* tachyzoites (RH) or with populations of RH that had been transfected with HA-tagging plasmids targeted to the indicated locus (RH::211460-3xHA, RH::204340-3xHA, and RH::PPM3C-3xHA). Lysates from infected HFFs were prepared and the HA-tagged proteins were detected by western blotting and probing with rat anti-HA antibodies. Rabbit anti-SAG2A staining was used as a loading control for total parasite protein. Approximate migration of a ladder of size standards (sizes in kDa) is indicated. B. Representative immunofluorescence microscopy images of endogenously tagged parasite proteins. The populations of endogenously tagged parasites analyzed in Fig. 2A were allowed to infect HFFs for 16 hours before the infected monolayers were fixed with methanol. The corresponding tagged proteins in parasites that had successfully incorporated the HA-tag were detected with rat anti-HA antibodies, while all tachyzoites were detected with mouse anti-SAG1 antibodies and the entire monolayer was visualized with differential interference microscopy (DIC). The arrows indicate localization of the endogenously tagged proteins outside of the parasites and within the PV. Scale bar is 10µm.

Using the HA-tagged 211460, 204340, and PPM3C parasite populations, we next sought to determine the localization of these proteins in infected cells. Using SignalP software (v5.0), all three proteins have strongly predicted signal peptides although in the case of 211460, this is only true if translation starts at the fourth in-frame methionine (position 61) relative to the protein sequence predicted on ToxoDB (v45). The results (**Fig. 2B**) show a clear PV-like signal outside of the parasites in the 211460-3xHA, 204340-4xHA, and PPM3C-3xHA populations, including at the periphery of the PV. The PV-localization for 211460 is further confirmed in the independently generated clonal line (**Fig. S1B**). Thus, we conclude that 211460, 204340, and PPM3C are at least transiently localized to the *Toxoplasma* PV during infection. Furthermore, we also assessed the localization of these proteins within the parasites themselves. The results (**Fig. S2**) show that while PPM3C appears to be present throughout the parasite, 211460 and 204340 show a clear, punctate staining pattern that largely co-localizes with the dense granule protein GRA7, suggesting that these two proteins are also GRA proteins. We therefore designate *204340* as *GRA54* for its GRA-like localization, and *211460* as *MYR4*, for reasons described below.

To assess their potential involvement in GRA effector translocation, we attempted to generate knockouts of our candidate genes in a strain of *Toxoplasma* that constitutively expresses an HA-tagged version of the MYR1-dependent secreted effector protein GRA16, RHΔ*gra16*::GRA16-HA (“parental”). To do this, we co-transfected a CRISPR/Cas9 sgRNA plasmid that targets the first exon of the relevant gene along with a pTKO2-CAT-mCherry plasmid (CAT encodes the chloramphenicol-resistance gene, chloramphenicol acetyl transferase; **Fig. 3A**). Following selection with chloramphenicol, we cloned the populations by limiting dilution and confirmed disruptive integration of the vector by PCR with gene-specific primers. Using this strategy, we were able to disrupt the genomic loci of *MYR3*, *GRA44*, *GRA45*, *CST1*, *GRA54*, *MYR4*, *PPM3C*, and *GRA7* (**Fig. S3**). Despite several attempts, however, we were unable to generate a *GRA52* mutant. This gene may be essential as it has a very negative CRISPR fitness score of −3.96 (35). Given that the *MAF1* locus is expanded in *Toxoplasma*, with 4 copies in RH parasites (36), we chose not to attempt a CRISPR/Cas9 approach to knockout MAF1a, and instead utilized a previously generated strain in which the entire *MAF1* cluster (including MAF1a and MAF1b) is deleted (31).

**Figure 3.**
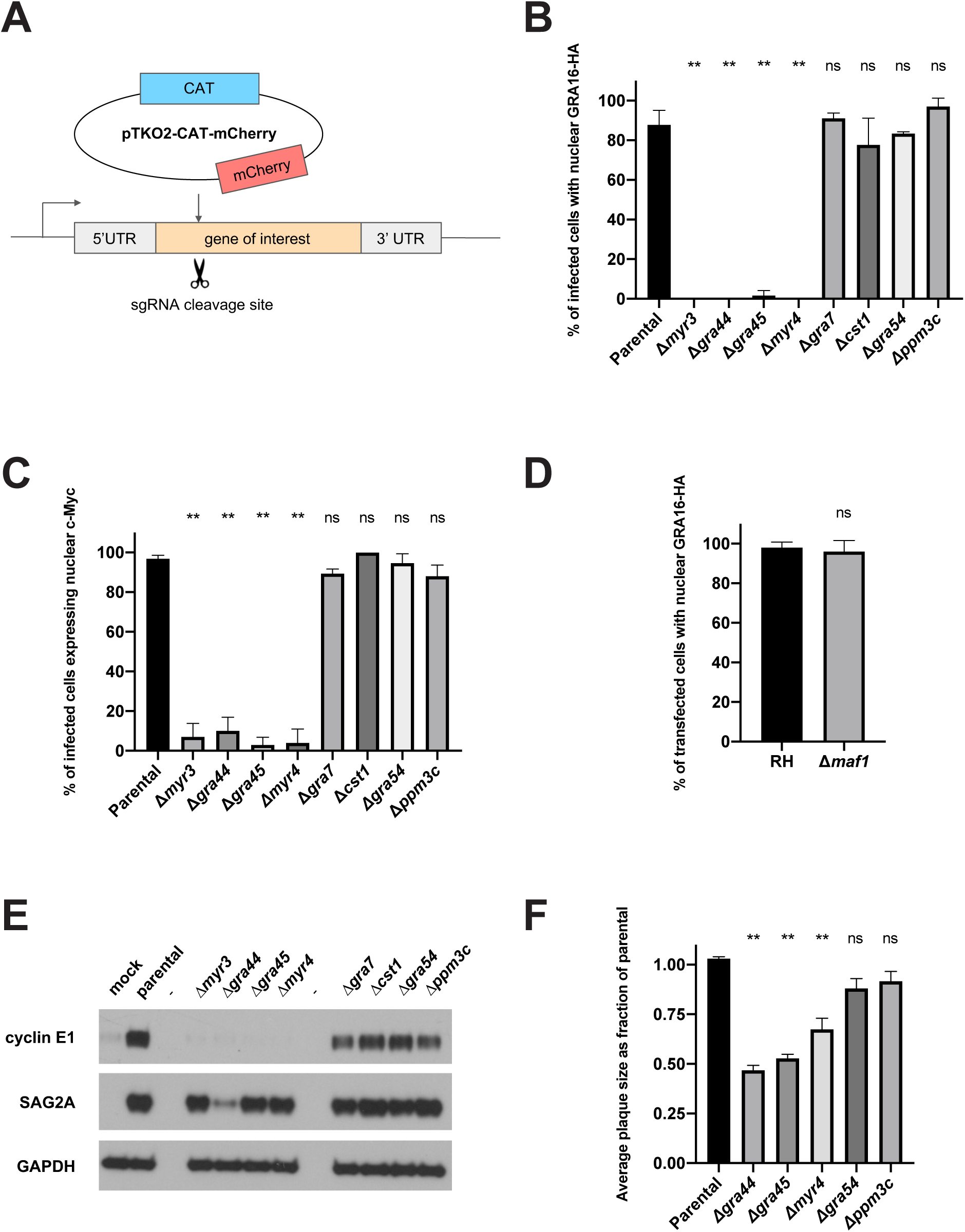
GRA44, GRA45, and MYR4 are required for *Toxoplasma* effector translocation and fully efficient growth *in vitro*. A. Schematic of CRISPR-mediated gene disruption of candidate genes followed by insertion of the pTKO2 plasmid carrying *mCherry* and a *chloramphenicol acetyl transferase* (*CAT*) gene for selection in chloramphenicol. B. Quantification of the percentage of infected cells showing GRA16-HA in the host nucleus via IFA. Tachyzoites were allowed to infect HFFs for 16 hours before the infected monolayers were fixed with methanol and stained with rat anti-HA antibodies. The averages are based on examination of at least 25 infected host cells per experiment from 2-5 biological replicates, and error bars indicate the standard deviation (SD). Statistics were performed using one-way ANOVA and Dunnett’s multiple comparisons test. ** indicates p<0.0001 and ns indicates nonsignificance (p>0.05) for the indicated strain relative to the parental control. C. Quantification of the percentage of infected cells showing upregulation of human c-Myc protein in the host nucleus via IFA. Tachyzoites were allowed to infect HFFs in serum-free media for 20 hours before the infected monolayers were fixed with methanol and stained with rabbit anti-c-Myc antibodies. Scoring and statistics are as for Fig. 3B. D. Quantification of the percentage of transfected, infected cells showing GRA16-HA in the host nucleus via IFA. Wild-type RHΔ*hpt* and RHΔ*maf1* tachyzoites were transiently transfected with a plasmid expressing GRA16-HA, and transfected parasites were allowed to infect HFFs for 16 hours before the infected monolayers were fixed with methanol and stained with rat anti-HA antibodies. The averages are based on the examination of 25 vacuoles from 2 biological replicates, and error bars indicate the SD. Statistics are as for Fig. 3B. E. Western blot of human cyclin E1 protein in infected cells. HFFs were infected with the indicated tachyzoites, or mock-treated with uninfected HFF lysate, for 18 hours before lysates were generated for immunoblotting. Lysates were analyzed by western blotting using mouse anti-cyclin E1 antibodies. Rabbit anti-SAG2A and mouse anti-GAPDH were used to assess the levels of parasite and host protein in the lysate respectively. “-“ indicates empty lanes. F. Quantification of plaque size. HFFs were infected with tachyzoites of the indicated strain for 7 days, fixed with methanol, and then stained with crystal violet. Plaque size was measured using ImageJ. Plaque areas were normalized to the median of the parental strain for each biological replicate. The averages are based on the results of at least 3 independent biological replicates, each with 2-3 technical replicates, and error bars represent the standard error of the mean. Statistics are as for Fig. 3B.

To determine if the absence of any of the candidate genes results in a defect in effector translocation across the PVM, we used IFA to assess both GRA16-HA export to the host nucleus and host c-Myc upregulation (which *Toxoplasma* induces during infection (37)) in the disrupted lines. Quantified results for all nine genes tested show that disruption of *GRA44*, *GRA45*, *MYR4* and the previously described *MYR3*, all resulted in a complete or near-complete block in GRA16 export to the host nucleus (**Figs. 3B, S4**) and a failure to upregulate host c-Myc (**Figs. 3C, S4**); on the other hand, disruption of *GRA7*, *CST1*, *GRA54*, or *PPM3C* resulted in no detectable effect on either of these two phenotypes. Additionally, we found that the previously generated Δ*maf1* strain also had normal GRA16 export to the host nucleus (**Fig. 3D**). These results indicate that of the nine genes tested here, only *MYR3*, *GRA44*, *GRA45* and *MYR4* are necessary for the translocation of GRA effectors across the PVM.

To test the generality of their role in effector translocation, we next assessed the impact of these gene disruptions on the upregulation of host cyclin E1 which has been shown to be dependent on export of the MYR1-dependent effector HCE1/TEEGR (6). The results showed that, as for GRA16, disruption of *MYR3*, *GRA44*, *GRA45*, and *MYR4* also resulted in a block in cyclin E1 upregulation in infected host cells, while no obvious defect was observed in the parasite lines disrupted in *GRA7*, *CST1, GRA54* and *PPM3C* (**Fig. 3E**). A repetition of the cyclin E1 western blot with higher parasite input reveals that the absence of cyclin E1 upregulation observed in Δ*gra44* parasites in Fig. 3E is not due to low parasite input in that particular experiment (**Fig. S5**). These results argue that *GRA44*, *GRA45* and *MYR4* are all required for translocation across the PVM of at least two independent GRA effectors.

Our previous work has shown that deletion of *MYR1*, *MYR2*, and *MYR3* results in a small but significant, negative effect on parasite growth *in vitro* (11). To determine if disruption of the three new genes involved in effector translocation described here has a similar impact, we infected HFF monolayers with each of the disrupted lines, fixed the monolayers 7 days post infection, and measured plaque size. The results show that the *Δmyr4, Δgra44* and *Δgra45* strains all exhibit a significant growth defect compared to the parental strain (**Fig. 3F**). We did not test for rescue of the growth phenotype with complementation due to limitations in selectable markers available in these strains. The *Δgra54* and *Δppm3c* strains, on the other hand, did not have significant growth defects, consistent with the growth defects observed being dependent on the respective genotype rather than nonspecific effects of the manipulations.

To confirm that ablation of *GRA44*, *GRA45*, and *MYR4* loci are responsible for the observed defect in GRA16 export, we transiently expressed a C-terminally V5-tagged version of each protein, driven by its native promoter, in the relevant disrupted line. These transiently transfected parasites were then assessed for GRA16-HA export to the host nucleus via IFA. The results showed that the parental and complemented strains had GRA16-HA signal in both the vacuole and host nucleus, while the parasites within the population that did not express the complementing transgene (as indicated by lack of anti-V5 staining) showed essentially no GRA16 in the host nucleus (**Figs. 4A, 4B**). Thus *GRA44*, *GRA45*, and *MYR4* are indeed essential for the translocation of effectors across the PVM, and we therefore designate *211460* as *MYR4,* consistent with previous nomenclature (10, 11).

**Figure 4.**
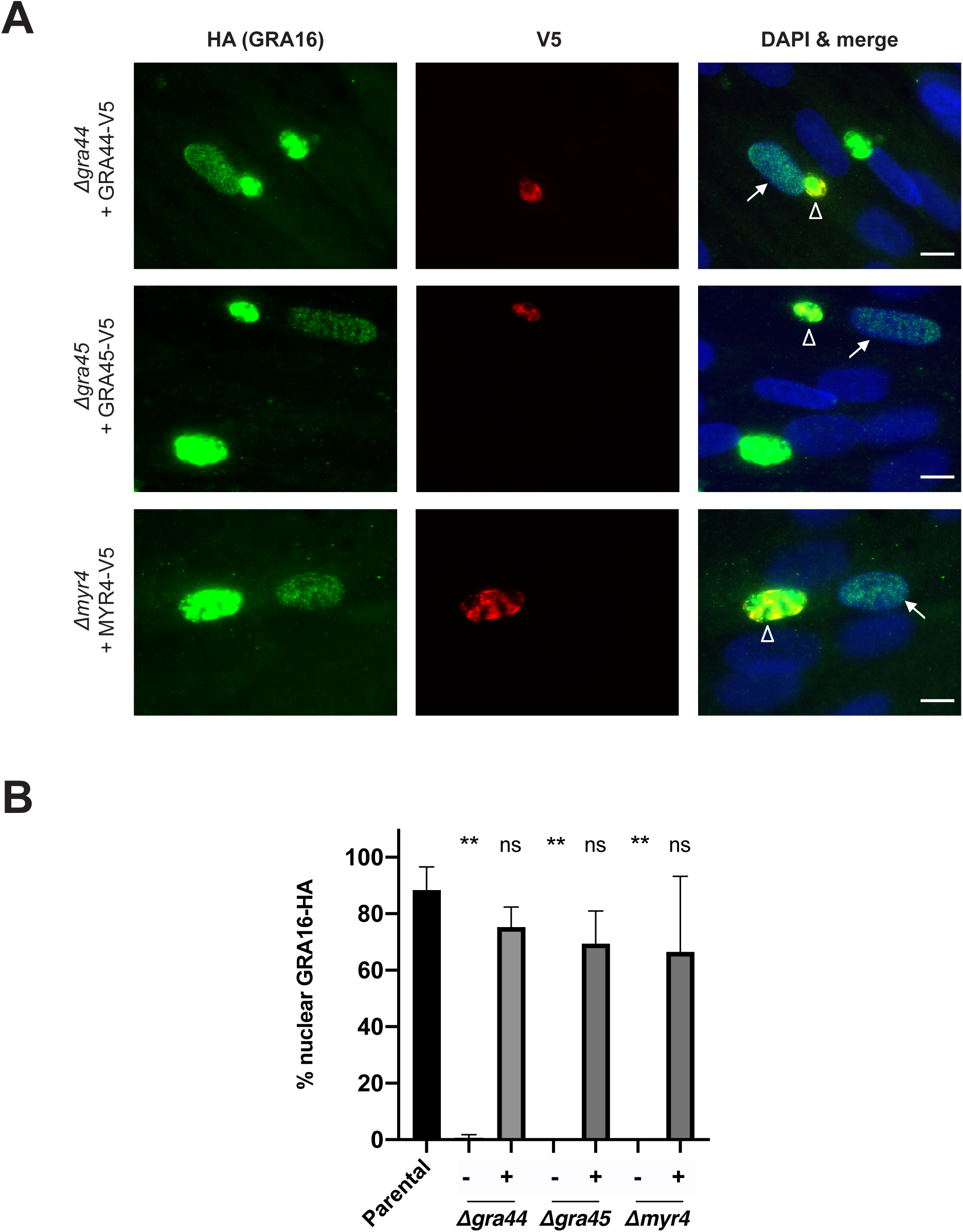
Ectopic protein expression rescues the effector translocation defect in Δ*gra44*, Δ*gra45*, and Δ*myr4* parasites. A. Representative immunofluorescence microscopy images of transiently expressed GRA44, GRA45, and MYR4 proteins. The indicated strains were transiently transfected with plasmids expressing the corresponding, C-terminally V5-tagged protein under its native promoter and the tachyzoites were allowed to infect HFFs for 18-22 hours before the infected monolayers were fixed with methanol. Localization of the V5-tagged proteins and rescue of the GRA16-HA host nuclear translocation were assessed by IFA using mouse anti-V5 and rat anti-HA antibodies, respectively. White arrows indicate a GRA16-HA positive host nucleus in a cell infected with tachyzoites expressing the indicated V5 tagged protein (white open arrowheads). Scale bar is 10µm. B. Quantification of the data represented in Fig. 4A showing the percentage of infected cells showing GRA16-HA in the host nucleus via IFA. The indicated strains were transiently transfected with either empty plasmid (-) or plasmids expressing the corresponding C-terminally V5-tagged protein (+) under its native promoter. Scoring and statistics are as for Fig. 3B, except for “+” conditions where only cells infected with V5-positive vacuoles were quantified.

Interestingly, GRA44, GRA45 and MYR4 all contain one or two instances of the three-amino-acid motif “RRL” (**Fig. 5A**), which has previously been shown to be the preferred sequence for cleavage by ASP5 protease (15). Indeed, cleavage at the three sites shown in GRA44 and GRA45 (27), as well as at the first “RRL” motif in the secreted GRA effector, GRA16 (15), has been experimentally confirmed. ASP5 is essential for the translocation of all GRA effectors so far tested (5, 6, 8, 9, 15–17) and it has previously been suggested that ASP5-mediated cleavage of some effectors is required to “license” them for translocation across the PVM, as appears to be the case in *Plasmodium* (38, 39). Given, however, that not all such effectors contain ASP5 processing motifs (e.g., GRA24 lacks the canonical “RRL” and shows no evidence of ASP5-dependent processing; (17)), and given that the three newly identified components of the translocation machinery identified here do, we hypothesized that ASP5’s essential contribution to effector translocation across the PVM might be in processing one or more components of the translocation machinery. We have previously shown that MYR1 is also processed by ASP5 at a “RRL” site but this does not appear to be required for MYR1 to function in effector translocation (24) and so we turned our attention to the newly identified translocation components identified here.

**Figure 5.**
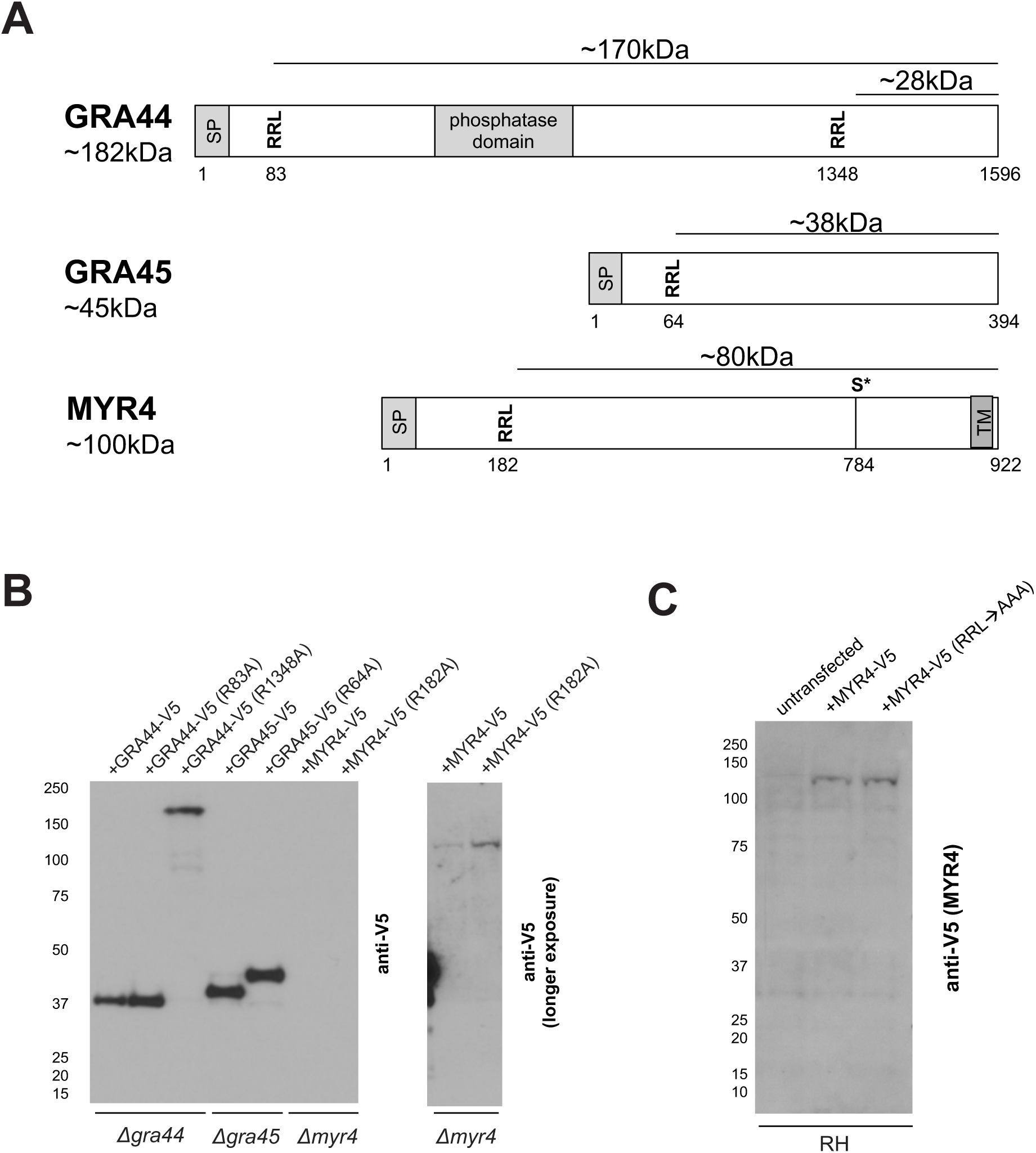
GRA44 and GRA45, but not MYR4, show evidence for processing at RRL sites. A. Schematic of GRA44, GRA45, and MYR4 protein sequence showing the location of predicted signal peptides (SP), RRL tripeptide sequences, previously identified phosphorylated serine residues (S*) and conserved domains, numbered in amino acid residues relative to the predicted N-terminus of the primary translation product. Approximate molecular weights (kDa) of the indicated portions are indicated. The amino acid sequence of MYR4 was determined using the 4th in-frame methionine relative to the protein predicted in ToxoDB (v45). Transmembrane domain prediction based on Phobius (Lukas Kall et al., Nucleic Acids Res 35:W429-32, 2007, https://doi.org/10.1093/nar/gkm256). B. Western blot of protein processing. The indicated parasite lines were transiently transfected with plasmids expressing C-terminally V5-tagged versions of either the indicated wildtype protein or a mutant version with the indicated RRL mutated to ARL (numbers indicate the amino acid position of the mutated arginine). These were then used to infect HFFs for 18 hours. Lysates were analyzed by western blotting using mouse anti-V5 antibodies to detect the C-terminally V5-tagged portions of each protein. Approximate migration of a ladder of size standards (sizes in kDa) is indicated. The right panel is a longer exposure of the right-most two lanes of the left panel. C. Western blot of MYR4 processing. RHΔ*hpt* (RH) parasites were transiently transfected with either WT or an RRL→AAA mutated version of C-terminally V5-tagged MYR4 and allowed to infect HFFs for 24 hours before lysates were generated for immunoblotting. Lysates were analyzed by western blotting using mouse anti-V5 antibodies to detect MYR4. Approximate migration of a ladder of size standards (sizes in kDa) is indicated.

To determine if processing at the “RRL” sites of GRA44, GRA45, and MYR4 is required for protein translocation activity, we mutated the ASP5 cleavage sites by converting the first arginine to an alanine (i.e., RRL→ARL) in the V5-tagged complementation plasmids for each gene, and transiently transfected these into the corresponding disrupted line. Western blots were then used to show that processing of GRA45 at its lone “RRL” and of GRA44 at its second “RRL” is indeed abrogated by the mutations (**Fig. 5B**). For the more N-terminal site in GRA44 (R83A), we cannot definitively confirm that the mutation abrogates ASP5 processing because GRA44 is epitope-tagged at its C-terminus and so, assuming cleavage at the two sites is an independent event, cleavage at the downstream site will produce a C-terminal, V5-tagged fragment whether or not cleavage occurs at R83A. We fully expect, however, that the RRL→ARL change disrupts ASP5 cleavage at this site because it did in the two other examples shown here (GRA45 and the downstream site in GRA44, R1348A) and because RRL→ARL mutations have previously been shown to disrupt the ASP5-dependent cleavage of other proteins (24, 27).

Interestingly, mutation of the “RRL” to an “ARL” in MYR4 did not appear to affect the processing of the protein (**Fig. 5B**). To rule out whether this is due to incomplete ablation of the ASP5 processing site with a single amino acid substitution, we assessed the processing of an RRL→AAA MYR4 mutant where the entire ASP5-processing motif is mutated to alanines. The results (**Fig. 5C**) show that the higher molecular weight product of MYR4 (∼130kD) does not change in mobility upon mutation of the entire “RRL” motif, and thus we conclude that little if any MYR4 is processed by ASP5. Note that, despite repeated attempts with large amounts of DNA, the signal for the transiently expressed MYR4 was never strong enough to confidently conclude whether a small amount of a processed form might be present in these transiently transfected parasites; we therefore cannot comment on whether the low-intensity, smaller molecular weight product of MYR4 (∼100kD) seen in long exposures of endogenously tagged wild type MYR4 (**Figs. 2A, S1A**) is a result of an ASP5 processing event.

Having generated the four RRL→ARL mutants, and having validated that ASP5 cleavage is ablated in at least two instances, we next tested each for its impact on the localization of the epitope-tagged, C-terminal portion of the protein and on the ability of the uncleaved protein to function; i.e., whether it can rescue the defect in effector protein translocation. The results show that the RRL→ARL mutated versions of each protein are still secreted into the PV, similar to the wild type copy (**Figs. 6A, 4A**), and are all able to rescue the translocation defect to a similar extent as the corresponding control (WT) plasmid (**Fig. 6B**). While the GRA45 R64A mutant did substantially rescue translocation, it did not consistently rescue to wildtype levels. Nevertheless, these data suggest that mutation of the “RRL” sites in GRA44, GRA45, and MYR4 to “ARL” does not substantially affect their function in effector protein translocation.

**Figure 6.**
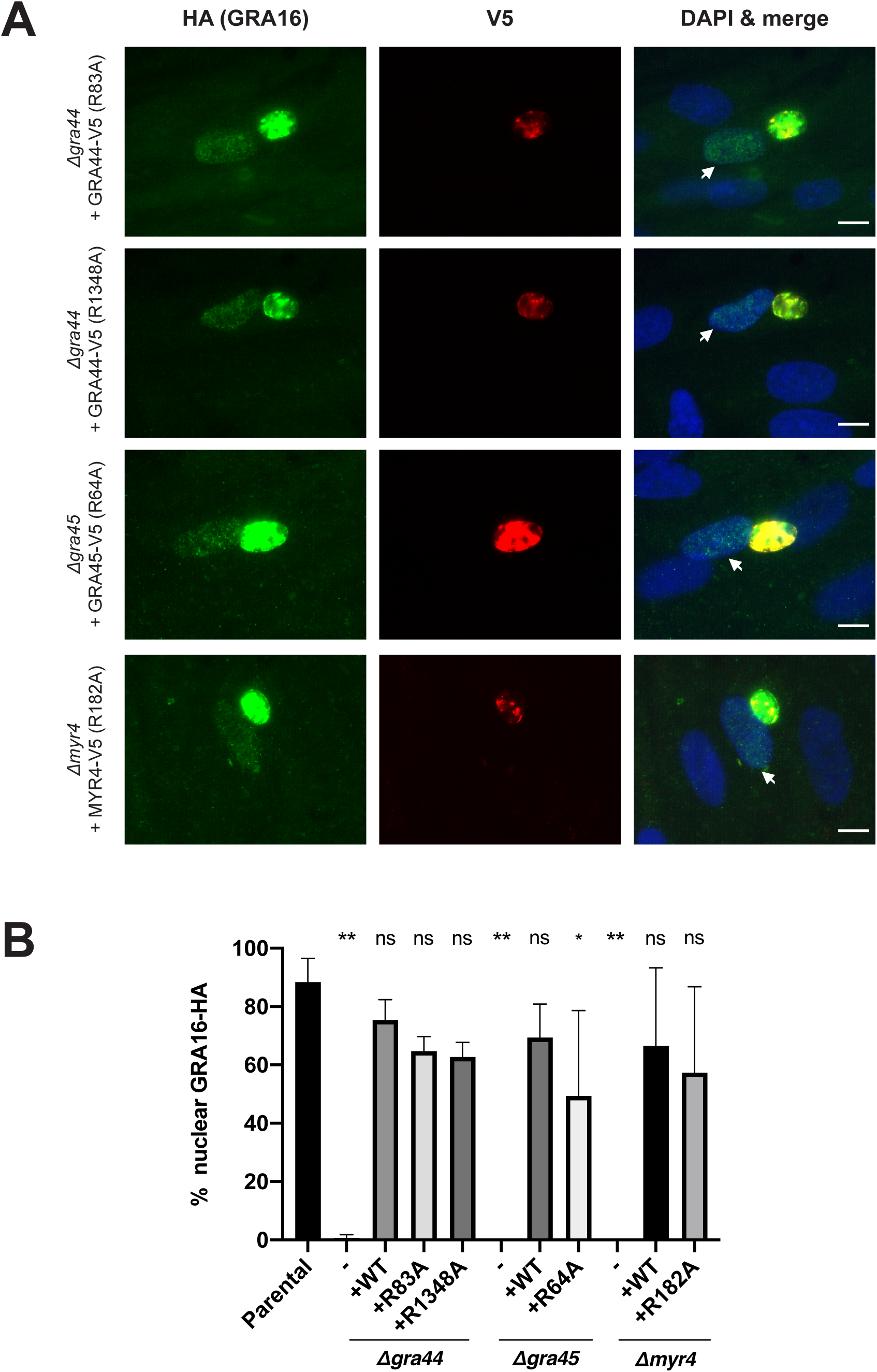
Ectopic expression of RRL mutants of GRA44, GRA45, and MYR4 rescues the translocation defect in Δ*gra44*, Δ*gra45*, and Δ*myr4* parasites. A. Representative immunofluorescence microscopy images of transiently expressed GRA44, GRA45, and MYR4 RRL→ARL mutated proteins. The indicated strains were transiently transfected with plasmids expressing the corresponding C-terminally V5-tagged protein under its native promoter and the tachyzoites were allowed to infect HFFs for 18-22 hours before the infected monolayers were fixed with methanol. Localization of the V5-tagged proteins and rescue of the GRA16-HA host nuclear translocation were assessed by IFA using mouse anti-V5 and rat anti-HA antibodies, respectively. White arrows indicate a GRA16-HA positive host nucleus in a cell infected with tachyzoites expressing the indicated V5 tagged protein. Scale bar is 10µm. B. Quantification of the data represented in Fig. 6A showing the percentage of infected cells showing GRA16-HA in the host nucleus via IFA. The indicated strains were transiently transfected with either empty plasmid (-) or plasmids expressing the corresponding C-terminally V5-tagged protein (+) under its native promoter. The data for the untransfected parental strain, the empty plasmid transfected strains, and the wildtype protein transfected strains are the same as in Fig. 4B and are included here for ease of comparison. Scoring and statistics are as for Fig. 4B except * indicates p=0.017.

## Discussion

Using affinity purification of MYR1 under conditions expected to retain associating partners, we identify three novel parasite proteins, GRA44, GRA45, and MYR4, as essential for the export of GRA effectors into infected cells. Additionally, we localize MYR4, as well as two additional MYR1-associating proteins, GRA54 and PPM3C, to the PV in infected cells. Altogether, eight proteins are now known to be necessary for effector export: the 3 described here and MYR1, MYR2, MYR3, ROP17 and ASP5 – summarized in Table S1 (10–12, 15–17). Besides ASP5, which localizes to the Golgi (15–17), these proteins all localize to the PV/PVM.

The newly identified components described here do not display any homology to known protein translocation machinery based on BLAST analysis results (BLASTP 2.10.0+), making it difficult to infer their functions and thus which, if any, are part of an actual translocon remains unknown. In addition to lacking homology to known translocation machinery, MYR4 and GRA45 do not have detectable homology to any other known, functional protein domains and neither do they share homology to proteins in any species outside of *Coccidia/Eimeriorina*. Like MYR1, MYR2, MYR3, and ROP17, however, MYR4, GRA44 and GRA45 all have clear orthologs in *Hammondia hammondi* and *Neospora caninum* (**Table S1**).

GRA44, by contrast, contains a putative phosphatase domain that shares homology to a region of the *Plasmodium* serine/threonine phosphatase UIS2 (28% identity over 21% of the protein; BLASTP 2.10.0+), which has recently been shown to localize to the *Plasmodium* PVM in liver stage parasites (40). Whether UIS2 plays a role in protein translocation in *Plasmodium* remains to be determined but this would be surprising given that none of the other components of the complex known to promote translocation in *Plasmodium* (known as PTEX) so far studied play a role in translocation in *Toxoplasma* (2). Additionally, whether this phosphatase domain is important for effector export in *Toxoplasma* is not yet known. Given that the kinase domain of ROP17 is necessary for GRA16 export (12) it is intriguing that two of the eight factors necessary for effector export are either a kinase or a phosphatase. There are numerous serine residues that are phosphorylated among MYR1, MYR2, MYR3, and MYR4, supporting the possibility that phosphorylation of the translocation machinery is critical to regulating its function in effector export. While this work was in progress, we learned of similar studies by Blakely, Arrizabalaga and colleagues who also found that GRA44 associates with MYR1 and is necessary for efficient c-Myc upregulation during infection (see accompanying manuscript). These latter authors used a knockdown approach to study GRA44 and saw a more dramatic impact of GRA44 loss on parasite growth than we report here for the GRA44 knockout; this might indicate that compensatory changes were selected for during the prolonged selection necessary to generate and expand our knockout clone, as was reported for AMA1 knockouts that showed dramatic up-regulation of the paralogue, AMA2 (41). Thus, transcriptomic analysis of the GRA44 knockout may reveal clues to its specific role(s) in *Toxoplasma* tachyzoites.

Our results expand the enigmas of why some parasite proteins are proteolytically processed by ASP5, and why ASP5 is essential for effector translocation across the PVM. MYR1, GRA44, and GRA45 all possess “RRL” motifs that appear to be cleaved in an ASP5-dependent manner yet, surprisingly, their function in the export of GRA16, and of GRA24 in the case of MYR1 (24), appears agnostic to mutation of these sites. For MYR1, we previously showed that the two domains generated by ASP5 processing stay connected through a disulfide bond after cleavage (11); it remains to be determined whether the polypeptides formed by RRL cleavage in GRA44 and GRA45 likewise associate in a similar manner. It is also important to note that our assays may not be sensitive enough to detect small changes in protein abundance in the host nucleus, and that it is the combination of multiple proteins not being processed by ASP5 that is deleterious to export in *Δasp5* mutants, rather than the result of failure to cleave any single protein.

Interestingly, there was a large number of proteins that were more highly enriched than MYR3 in our immunoprecipitations with MYR1, and it remains a strong possibility that additional MYR1-associating proteins are involved in effector translocation. Due to the large number of enriched proteins and the limited throughput of our approach, we were unable to investigate all candidates for such a role; nevertheless, our data showing that GRA16-HA export is not lost in parasites disrupted for *GRA7*, *CST1*, *MAF1*, *PPM3C*, or *GRA54* strongly suggests that it is not general PV/PVM disruption that results in the loss of effector translocation. Further work will be needed to determine which of the remaining proteins we see enriched in the MYR1 immunoprecipitations are there because of specific association with MYR1 vs. nonspecific associations of proteins within the PV/PVM due to association within lipid rafts or other entities.

Lastly, none of *GRA44*, *GRA45*, or *MYR4* were identified in the forward genetic screen of parasites that are unable to induce c-Myc (10). This could be due to the growth defects observed in *Δmyr4, Δgra44,* and Δ*gra45* parasites shown here since parasites with null mutations in these genes might be lost during the 7-8 rounds of selection used in that screen due to a fitness disadvantage. Alternatively, the mutagenesis-based genetic screen was not saturating and so a more comprehensive, genome-wide screen using CRISPR/Cas9 technologies might reveal these and other genes responsible for effector translocation in *Toxoplasma*. Regardless, our finding of three new components of the export machinery provides a richer understanding of how *Toxoplasma* delivers effectors into host cells. Future work will determine the precise function of each, including how they interact, the role of ASP5 cleavage, and which, if any, constitutes the actual translocon.

## Materials and Methods

### Parasite strains, culture and infections

All *Toxoplasma* tachyzoites used in this study are in the Type I “RH” background, either RH::*MYR1-3xHA* (11), RHΔ*gra16::GRA16HA* (6), RHΔ*maf1* (31), RHΔ*hpt* (42), or RHΔ*hpt*Δ*ku80* (43). These tachyzoites, and all subsequently generated lines, were propagated in human foreskin fibroblasts (HFFs) cultured in complete Dulbecco’s Modified Eagle Medium (cDMEM) supplemented with 10% heat-inactivated fetal bovine serum (FBS; HyClone, Logan, UT), 2 mM L-glutamine, 100 U/ml penicillin and 100 µg/ml streptomycin at 37 °C with 5% CO_2_. The HFFs were obtained from the neonatal clinic at Stanford University following routine circumcisions that are performed at the request of the parents for cultural, health or other personal medical reasons (i.e., not in any way related to research). These foreskins, which would otherwise be discarded, are fully de-identified and therefore do not constitute “human subjects research”.

Prior to infection, parasites were scraped and syringe-lysed using a 27 G needle, counted using a hemocytometer, and added to HFFs. “Mock” infection was done by first syringe-lysing uninfected HFFs, processing this in the same manner as done for the infected cells, and then adding the same volume of the resulting material as used for infections. For experiments where human c-Myc protein was detected, the parasites were added to HFFs in media containing 0% serum.

### Immunofluorescence assay (IFA)

Infected cells grown on glass coverslips were fixed and permeabilized using 100% cold methanol for 10 min. Samples were washed 3x with PBS and blocked using 3% BSA in PBS for 1 hour at room temperature (RT). HA was detected with rat monoclonal anti-HA antibody 3F10 (Roche), SAG1 was detected with mouse anti-SAG1 monoclonal antibody DG52 (44), GRA7 was detected with rabbit anti-GRA7 antibodies (45), V5 was detected with mouse anti-V5 tag monoclonal antibody (Invitrogen), and c-Myc was detected with rabbit monoclonal anti-c-Myc antibody Y69 (Abcam). Primary antibodies were detected with goat polyclonal Alexa Fluor-conjugated secondary antibodies (Invitrogen). Primary and secondary antibodies were both diluted in 3% BSA in PBS. Coverslips were incubated with primary antibodies for 1 hour at RT, washed, and incubated with secondary antibodies for 1 hour at RT. Vectashield with DAPI stain (Vector Laboratories) was used to mount the coverslips on slides. Fluorescence was detected using wide-field epifluorescence microscopy and images were analyzed using ImageJ. All images shown for any given condition/staining in any given comparison/dataset were obtained using identical parameters.

### Transfections

All transfections were performed using the Amaxa 4D Nucleofector (Lonza) model. Tachyzoites were mechanically released in PBS, pelleted, and resuspended in 20 µL P3 Primary Cell Nucleofector Solution (Lonza) with 5-25 µg DNA for transfection. After transfection, parasites were allowed to infect HFFs in DMEM.

### Plasmid construction

For gene disruption plasmids: gRNAs designed against a PAM site of each gene of interest were cloned into the pU6-Universal plasmid. pU6-Universal was a gift from Sebastian Lourido (Addgene plasmid # 52694; http://n2t.net/addgene:52694; RRID:Addgene_52694).

For ectopic expression plasmids: The pGRA-V5 plasmid was created by replacing the HA tag sequence in the pGRA-HPT-HA plasmid (46) with the V5 tag DNA sequence (GGCAAGCCCATCCCCAACCCCCTGCTGGGCCTGGACAGCAC) and removing the HPT resistance cassette using standard molecular biology techniques. The pX-V5 plasmid was created by removing the GRA1 promoter from pGRA-V5 using standard molecular biology techniques. Complementation plasmids to ectopically express V5 tagged proteins off their native promoters were created by PCR amplification of the open reading frame of each gene, minus the stop codon, plus ∼2000 bp upstream of the start codon to include the native promoter, followed by cloning into pX-V5 using Gibson Assembly (NEB). RRL→ARL or RRL→AAA mutated complementation plasmids were generated using overlap extension PCR using primers harboring the mutation and cloning the resultant products into pX-V5 using Gibson Assembly (NEB).

For endogenous tagging plasmids: Approximately 1500-3000 bp of the 3’ coding sequence of each gene was amplified from RH genomic DNA and cloned into the pTKO2-HPT-3xHA plasmid (11) using either Gibson Assembly (NEB) or by cloning into the EcoRV and NotI restriction sites.

A list of all primers and plasmids used and generated in this study can be found in **File S2**.

### Endogenous tagging

Endogenous tagging plasmids were transfected into *Toxoplasma* via electroporation. Tachyzoites were allowed to infect HFFs in T25 flasks for 24 hours, after which the medium was changed to complete DMEM supplemented with 50 μg/ml mycophenolic acid and 50 μg/ml xanthine for selection for the hypoxanthine-xanthine-guanine-phosphoribosyltransferae (HXGPRT or HPT) marker for 3-5 days.

### Gene disruption

A list of all sgRNA sequences used in this study can be found in **File S2**. RHΔ*gra16*::GRA16HA tachyzoites were transfected with pTKO2-CAT-mCherry (CAT is chloramphenicol acetyl transferase which confers resistance to chloramphenicol; the plasmid was a gift from Ian Foe and Matthew Bogyo (47)) and the corresponding modified pU6-sgRNA plasmid and allowed to infect HFFs for 24-48 hours. For gene disruption of MYR3, the previously published pSAG1:U6-Cas9:sgMYR3 plasmid was used instead (11). Between 24-48 hours after transfection, DMEM media with 80 µM chloramphenicol was added to the cells. The media was replaced with fresh chloramphenicol-supplemented media every 48-72 hours. After at least 7 days in selection, single clones were selected from the transfected populations in 96 well plates using limiting dilution. Single clones were maintained in chloramphenicol-supplemented media until confirmation of the genetic disruption.

### Ectopic expression

Plasmids for ectopic expression were transiently transfected into *Toxoplasma* using electroporation. Tachyzoites were allowed to infect HFFs for 18-24 hours before assessing for expression of the ectopically expressed protein via either IFA or western blotting.

### Western blotting

Cell lysates were prepared at the indicated time points post-infection in Laemmli sample buffer (BioRad). The samples were boiled for 5 min, separated by SDS-PAGE, and transferred to polyvinylidene difluoride (PVDF) membranes. Membranes were blocked with 5% nonfat dry milk in TBS supplemented with 0.5% Tween-20, and proteins were detected by incubation with primary antibodies diluted in blocking buffer followed by incubation with secondary antibodies (raised in goat against the appropriate species) conjugated to horseradish peroxidase (HRP) and diluted in blocking buffer. HA was detected using a horseradish peroxidase (HRP)-conjugated HA antibody (Roche cat no. 12013819001), SAG2A was detected using rabbit polyclonal anti-SAG2A antibodies (48), Cyclin E1 was detected using mouse monoclonal antibody HE12 (Santa Cruz Biotechnology), and GAPDH was detected using mouse monoclonal anti-GAPDH antibody 6C5 (Calbiochem). Horseradish peroxidase (HRP) was detected using enhanced chemiluminescence (ECL) kit (Pierce).

### Plaque assay

Parasites were syringe-released from HFFs and added to confluent HFFs in T25 flasks. After 7 days, the infected monolayers were washed with PBS, fixed with methanol, and stained with crystal violet. Plaque area was measured using ImageJ.

### Immunoprecipitations (IPs) for mass spectrometry

IPs to identify MYR1-interacting proteins in HFFs were performed as follows. One 15-cm dish of HFFs for each infection condition was grown to confluence. HFFs were infected with either 15 x 10^6^ RH::*MYR1-3xHA* or RHΔ*hpt* parasites for 24 hours. Infected cells were washed 3 times in cold PBS and then scraped into 1 mL cold cell lysis buffer (50mM Tris (pH 8.0), 150mM NaCl, 0.1% (v/v) Nonidet P-40 Alternative [CAS no. 9016-45-9]) supplemented with complete protease inhibitor cocktail (cOmplete, EDTA-free [Roche]). Cell lysates were passed 3 times through a 25 G needle, followed by 3 times through a 27 G needle, followed by sonication on ice (Branson Sonifier 250), with 3 pulses of 10 s at 50% duty cycle and output control 2. Cell lysates were spun at 1000 × *g* for 10 min to remove insoluble material and unlysed cells. Lysates were added to 100 µL magnetic beads conjugated to anti-HA antibodies (Pierce) and incubated overnight rotating at 4 °C. Unbound protein lysate was removed, and the anti-HA magnetic beads were then washed 10 times in cell lysis buffer. HA-tagged MYR1, and associated proteins, were eluted in 60 µL pH 2.0 buffer (Pierce) for 10 min at 50 °C to dissociate proteins from the antibody-conjugated beads. The elutions were immediately neutralized 1:10 with pH 8.5 neutralization buffer (Pierce).

### Mass spectrometry sample preparation

45 µL of each IP elution was combined with 15 µL of 4X Laemmli sample buffer supplemented with BME (BioRad), boiled for 10 min at 95 °C, and loaded on a Bolt 4-12% Bis-Tris gel (Invitrogen). The samples were resolved for approximately 8 min at 150V. The gel was washed once in UltraPure water (Thermo), fixed in 50% methanol and 7% acetic acid for 15 min, followed by 3 additional washes with UltraPure water. The gel was stained for 10 min with GelCode Blue (Thermo) and washed with UltraPure water for an additional 20 min. One gel band (approx. 1.5 cm in size) for each condition was excised and de-stained for 2 hours in a 50% methanol and 10% acetic acid solution, followed by a 30 min soak in UltraPure water. Each gel slice was cut into 1 mm x 1 mm squares, covered in 1% acetic acid solution, and stored at 4 °C until the in-gel digestion could be performed.

To prepare samples for mass spectrometry, the 1% acetic acid solution was removed, 10 µl of 50 mM DTT was added, and volume was increased to 100 µl with 50 mM ammonium bicarbonate. Samples were incubated at 55 °C for 30 min. Samples were then brought down to RT, DTT solution was removed, 10 µl of 100 mM acrylamide (propionamide) was added and volume was again normalized to 100 µl with 50 mM ammonium bicarbonate followed by an incubation at RT for 30 min. Acrylamide solution was removed, 10 µl (0.125 µg) of Trypsin/LysC (Promega) solution was added and another 50 µl of 50 mM ammonium bicarbonate was added to cover the gel pieces. Samples were incubated overnight at 37 °C for peptide digestion. Solution consisting of digested peptides was collected in fresh Eppendorf tubes, 50 µl of extraction buffer (70% acetonitrile, 29% water, 1% formic acid) was added to gel pieces, incubated at 37 °C for 10 min, centrifuged at 10,000 x g for 2 minutes and collected in the same tubes consisting of previous elute. This extraction was repeated one more time. Collected extracted peptides were dried to completion in a speed vac and stored at 4 °C until ready for mass spectrometry.

### Mass spectrometry

Eluted, dried peptides were dissolved in 12.5 μl of 2% acetonitrile and 0.1% formic acid and 3 μl was injected into an in-house packed C18 reversed phase analytical column (15 cm in length). Peptides were separated using a Waters M-Class UPLC, operated at 450 nL/min using a linear 80 minute gradient from 4-40% mobile phase B. Mobile phase A consisted of 0.2% formic acid, 99.8% water, Mobile phase B was 0.2% formic acid, 99.8% acetonitrile. Ions were detected using an Orbitrap Fusion mass spectrometer operating in a data dependent fashion using typical “top speed” methodologies. Ions were selected for fragmentation based on the most intense multiply charged precursor ions using Collision induced dissociation (CID). Data from these analyses was then transferred for analysis.

### Mass spectrometric analysis

The .RAW data were searched using MaxQuant version 1.6.1.0 against the canonical human database from UniProt, *Toxoplasma* GT1 databases from ToxoDB (versions 7.3 and 37.0), and the built-in contaminant database. Specific parameters used in the MaxQuant analysis can be found in **File S1**. Peptide and protein identifications were filtered to a 1% false discovery rate (FDR) and reversed proteins, contaminants, and proteins only identified by a single modification site, were removed from the dataset. MYR1-3xHA enrichment over the non-HA tagged RH was determined by adding 1 to each spectral count (tandem MS [MS/MS count]) and calculating the NSAF (number of spectral counts identifying a protein divided by the protein’s length, divided by the sum of all spectral counts/lengths for all proteins in the experiment). The average MYR1-3xHA enrichment from the two biological replicates (IP 1 and IP 2) was used to determine the protein ranking.

### Data availability

The mass spectrometry proteomics data have been deposited to the ProteomeXchange Consortium (http://proteomecentral.proteomexchange.org) via the PRIDE partner repository (49) with the dataset identifier PXD016383.

## Acknowledgements

We thank all members of our laboratory, as well as Melanie Espiritu for help with tissue culture and ordering, Ian Foe and Matthew Bogyo for providing reagents, Ryan Leib and Kratika Singhal for helpful advice regarding mass spectrometry, and Will Blakely and Gustavo Arrizabalaga for helpful discussions and exchange of data prior to publication. Special thanks to the Vincent Coates Foundation Mass Spectrometry Laboratory at Stanford University Mass Spectrometry (SUMS) for assistance in processing mass spectrometry samples.

This project has been funded in whole or part with federal funds from the U.S. National Institute of Allergy and Infectious Diseases, National Institutes of Health, Department of Health and Human Services, under awards NIH RO1-AI021423 (J.C.B.) and NIH RO1-AI129529 (J.C.B.), NIH T32-AI732832 (A.M.C.), NIH T32-AI007328 (T.C.T.), NIH T32-AI732832 (A.G.M.), NIH F31-AI120649 (N.D.M.), with funds from the National Science Foundation Graduate Research Fellowship Program (https://www.nsfgrfp.org/) under grant DGE-114747 (A.M.C.), with support from the Stanford Bio-X Graduate Research Fellowship (T.C.T.), and with grant NIH P30 CA124435 for utilization of the Stanford Cancer Institute Proteomics/Mass Spectrometry Shared Resource. The funders had no role in study design, data collection and analysis, decision to publish, or preparation of the manuscript.

## Figures and Figure Legends

**Supplemental File 1.**

Mass spectrometry analysis parameters and results for proteins that coimmunoprecipitate with MYR1-3xHA-expressing and untagged RH parasites. For all sheets, the IDs corresponding to the majority proteins, i.e., the proteins which contained at least half of the peptides belonging to a protein group (grouping of proteins which cannot be unambiguously identified by unique peptides), the number of spectral counts (MS/MS count), the average NSAF enrichment score (MYR1/RH Enrichment, as further elaborated in Materials and Methods), and the protein rank as defined by the enrichment score corresponding to each grouping are shown. The gene product (for *Toxoplasma* proteins) or associated gene name (for human proteins) for the first listed protein ID in each row is shown in the Description column. Sheet 1 (“Toxo_proteins”) shows the experimental data sets for *Toxoplasma* proteins only, listed in rank order by the average NSAF enrichment from both experiments. Sheet 2 (“All_proteins”) shows the experimental data sets for both human and *Toxoplasma* proteins, listed in rank order by the average NSAF enrichment from both experiments. Sheet 3 (“Parameters”) shows the parameters used in the MaxQuant analysis.

**Supplemental File 2.**

Primers, sgRNA sequences, and plasmids used and/or generated in this study.

**Supplemental Table 1.**

Summary of *Toxoplasma* genes necessary for effector translocation. The number of predicted transmembrane domains, number of “RRL” motifs, and CRISPR phenotype score are listed for each *Toxoplasma* gene necessary for effector translocation identified thus far. Additionally, the percent identities of each of these genes to their orthologs in *Hammondia hammondi* and *Neospora caninum*, and whether the “RRL” sequences are conserved in these species are also listed. Transmembrane domain prediction based on Phobius (Lukas Kall et al., Nucleic Acids Res 35:W429-32, 2007, https://doi.org/10.1093/nar/gkm256). CRISPR phenotype scores are from Sidik et al. (Cell 166(6):1423-1435.e12, https://10.1016/j.cell.2016.08.019). Identity calculated by comparison to head-to-head comparison of ortholog in indicated species using Sequence Manipulation Suite (Stothard P, Biotechniques 28:1102-1104, https://doi.org/10.2144/00286ir01).

**Supplemental Figure 1.**

A. Western blot of endogenously tagged 211460-3xHA single clone and population. HFFs were infected with RHΔ*hpt*Δ*ku80* tachyzoites (RH) or endogenously tagged RH::*211460-3xHA* parasites (either from the population or an independently generated single clone). Lysates from infected HFFs were prepared and 211460-3xHA was detected by western blotting using rat anti-HA antibodies. Rabbit anti-SAG2A staining was used as a loading control for total parasite protein. The western blot for the 211460-3xHA population is the same data as presented in Fig 2A. Approximate migration of a ladder of size standards (sizes in kDa) is indicated.

B. Immunofluorescence microscopy of endogenously tagged 211460-3xHA from an independently generated single clone. Tachyzoites were allowed to infect HFFs for 16 hours before the infected monolayer was fixed with methanol. 211460-3xHA was detected with rat anti-HA antibodies, *Toxoplasma* tachyzoites were detected with mouse anti-SAG1 antibodies, and the infected monolayer was visualized with DIC. Scale bar is 10µm.

**Supplemental Figure 2.**

Immunofluorescence microscopy of endogenously tagged proteins in extracellular parasites. The populations of endogenously tagged parasites analyzed in Fig. 2A were seeded onto empty coverslips before being fixed with methanol. The corresponding tagged proteins were detected with rat anti-HA antibodies, the marker for dense granule proteins, GRA7, was detected with rabbit anti-GRA7 antibodies, and the parasites were visualized with differential interference microscopy (DIC). Scale bar is 5µm.

**Supplemental Figure 3.**

A. Schematic of CRISPR-mediated gene disruption of candidate genes. Primers flanking the guide-targeted region, indicated by “Forward” and “Reverse”, were constructed to amplify a ∼1000bp region of the native, uninterrupted gene. pTKO2-CAT-mCherry is the plasmid used for integration and selection.

B. PCR amplifications of genomic DNA from RHΔ*gra16*::GRA16-HA parasites (parental) and from a chloramphenicol-resistant (CAT^+^) clonal strain with disruption of the indicated gene using the forward and reverse primers shown in Panel A. Sizes (base pairs) of the standard ladder are shown. Bands of the expected size in the parental strain (∼1000bp) and either lack of a band or presence of altered bands in the disrupted strains, indicate insertion of the selection plasmid within the targeted gene, as indicated (e.g., Δ*myr3* is a strain with a disruption of the *MYR3* locus).

**Supplemental Figure 4.**

Immunofluorescence microscopy of GRA16-HA nuclear localization and human nuclear c-Myc expression in HFFs infected with the indicated disrupted parasite strains. Tachyzoites were allowed to infect HFFs (without serum) for 18 hours before the infected monolayers were fixed with methanol and stained with rat anti-HA antibodies and rabbit anti-c-Myc antibodies. Host nuclei were visualized using DAPI. Scale bar is 20µm.

**Supplemental Figure 5.**

Western blot of human cyclin E1 protein in cells infected with the indicated parasite strain. HFFs were infected with the indicated strain of tachyzoites, or mock-treated with uninfected HFF lysate, for 20 hours before lysates were generated for immunoblotting. Lysates were analyzed by western blotting using mouse anti-cyclin E1 antibodies. Rabbit anti-SAG2A was used to assess the levels of parasite protein in the lysate.

