## Supplemental Figures for "Co-immunoprecipitation with MYR1 identifies three additional proteins within the *Toxoplasma* parasitophorous vacuole required for translocation of dense granule effectors into host cells"

| *Toxoplasma gondii* | | | | | | *Hammondia hammondi* | | *Neospora caninum* | | |
|---|---|---|---|---|---|---|---|---|---|---|
| Gene ID | Alias | Localization | Predicted TMs | Phenotype score | RRLs | Amino acid identity | RRL(s) conserved | Amino acid identity | RRL(s) conserved | Comments |
| 254470 | MYR1 | PV | 1 | 0.88 | 1 | 84% | 1 of 1 | 44% | 1 of 1 | A second RRL exists in middle of NcMYR1 |
| 270700 | MYR2 | PV | 2 | 2.39 | 0 | 84% | n/a | 36% | n/a | An RRL exists in the C-term of NcMYR2 |
| 237230 | MYR3 | PV | 1 | 2.83 | 0 | 72% | n/a | 31% | n/a | |
| 211460 | MYR4 | PV | 1 | 0.25 | 1 | 84% | 1 of 1 | 48% | 0 of 1 | |
| 228170 | GRA44 | PV | 0 | -3.28 | 2 | 94% | 2 of 2 | 68% | 2 of 2 | |
| 316250 | GRA45 | PV | 0 | 1.15 | 1 | 92% | 0 of 1 | 70% | 1 of 1 | An additional RRL exists in the C-term of NcGRA45 |
| 258580 | ROP17 | PV | 0 | 1.29 | 2 | 83% | 2 of 2 | 52% | 0 of 2 | |
| 242720 | ASP5 | Golgi | 3 | -1.45 | 1 | 90% | 0 of 1 | 66% | 0 of 1 | |

### Supplemental Table 1.

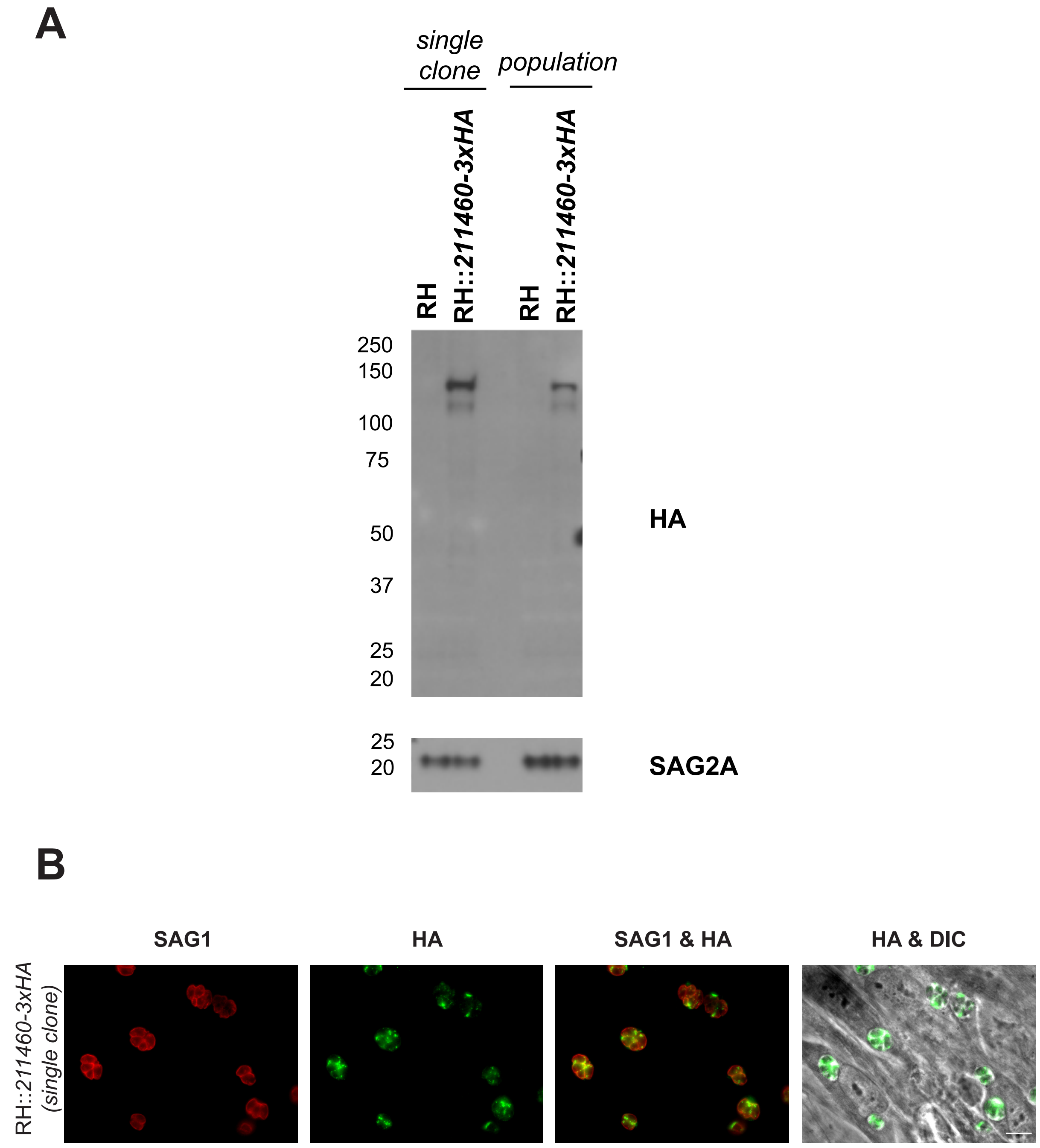


**Supplemental Figure 1.**

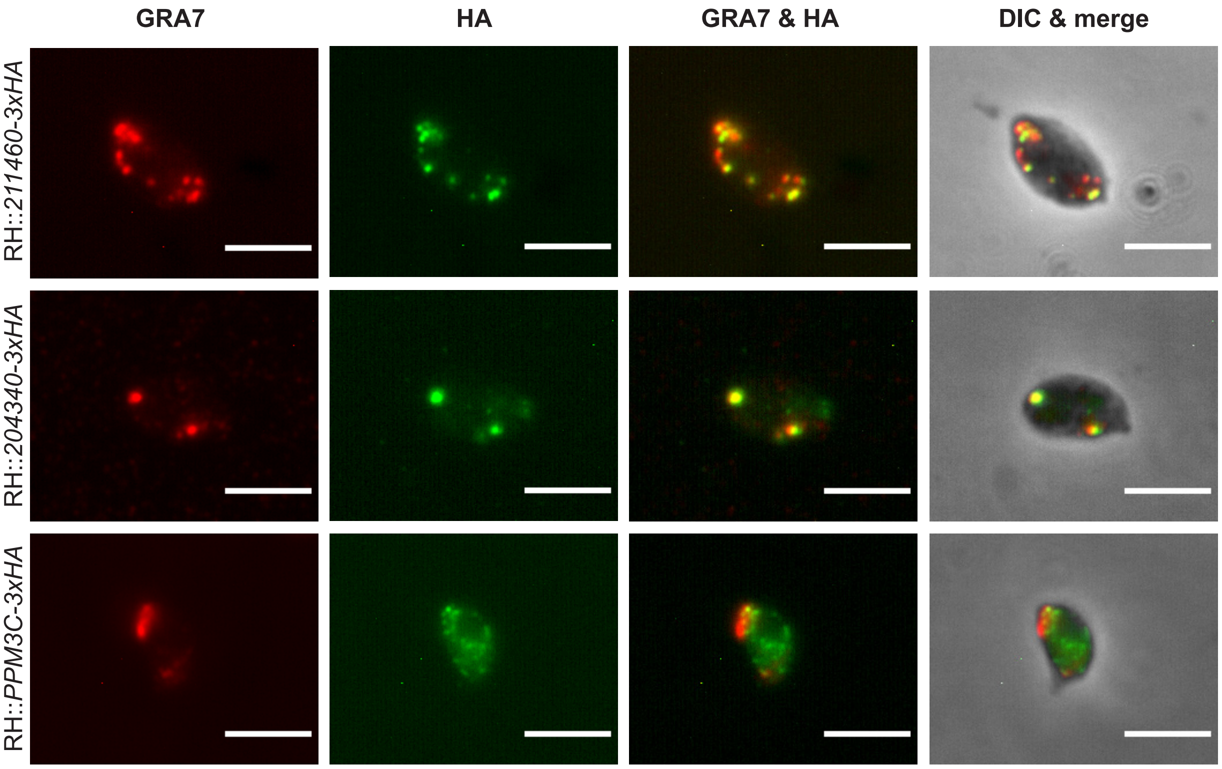


**Supplemental Figure 2.**

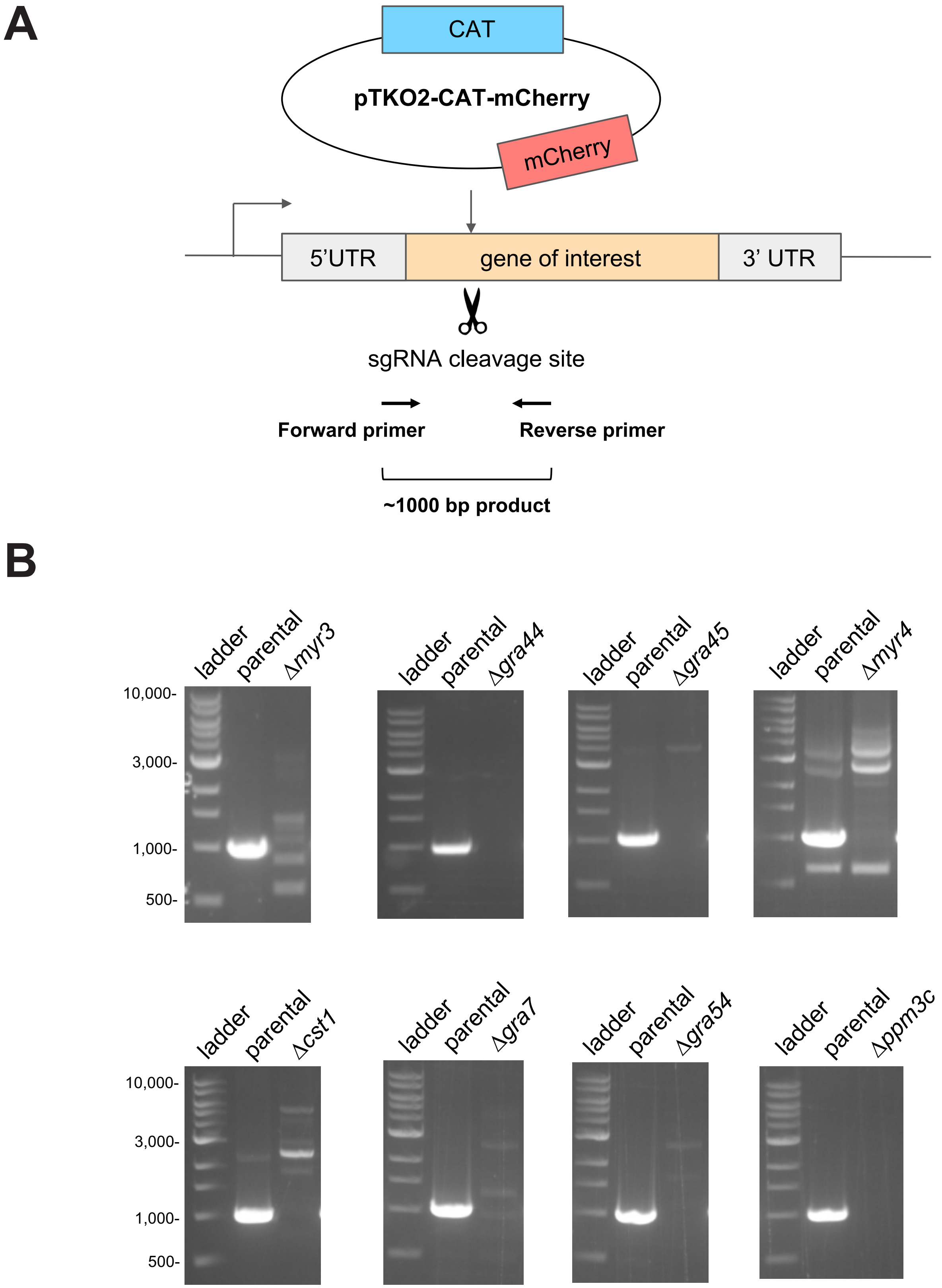


**Supplemental Figure 3.**

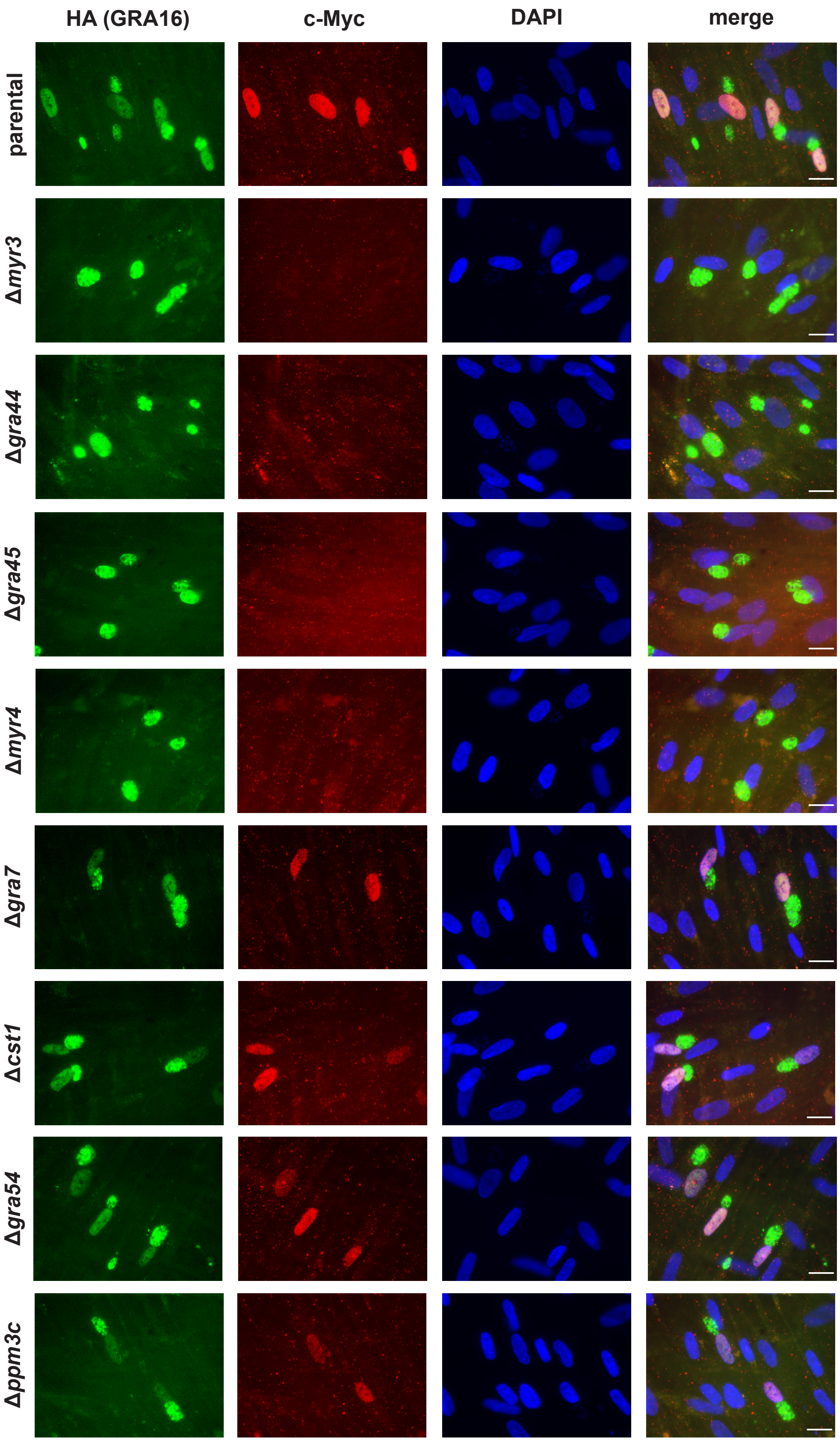


**Supplemental Figure 4.**

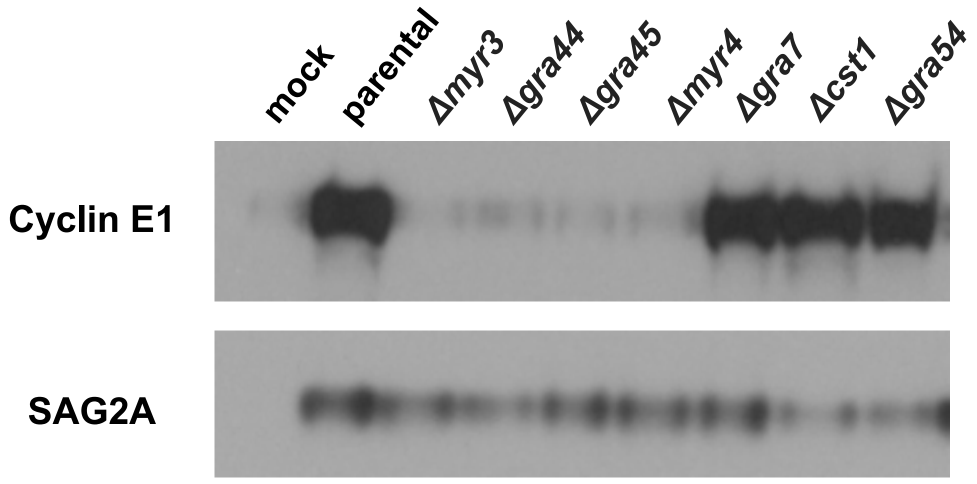


**Supplemental Figure 5.**
